# Chromatin remodeling integrates vertebrate body axis elongation and cell fate determination

**DOI:** 10.1101/2025.03.11.642544

**Authors:** Shukai Li, Viet Tien Nguyen, Wanjuan Bu, Hailang Fan, Deng Pan, Zixing Wang, Dongyan Chen, Dake Zhang, Yuliang Jin, Yubo Fan, Jing Du

**Affiliations:** Key Laboratory for Biomechanics and Mechanobiology of Ministry of Education, Beijing Advanced Innovation Center for Biomedical Engineering, National Medical Innovation Platform for Industry-Education Integration in Advanced Medical Devices (Interdiscipline of Medicine and Engineering), School of Biological Science and Medical Engineering, Beihang University, Beijing, 100190, China; Key Laboratory of Biomechanics and Mechanobiology, Ministry of Education; Key Laboratory of Innovation and Transformation of Advanced Medical Devices, Ministry of Industry and Information Technology; National Medical Innovation Platform for Industry-Education Integration in Advanced Medical Devices (Interdiscipline of Medicine and Engineering); School of Engineering Medicine, Beihang University, Beijing, 100191, China; Institute of Theoretical Physics, Chinese Academy of Sciences, Beijing 100190, China; Tianjin Key Laboratory of Tumor Microenvironment and Neurovascular Regulation, Department of Histology and Embryology, School of Medicine, Nankai University, Tianjin 300071, China

**Author notes:** These authors contributed equally (Shukai Li, Viet Tien Nguyen, Wanjuan Bu, Hailang Fan).

## Abstract

The effective elongation of the vertebrate body axis depends on the precise coordination of cell movements and cell fate determination within the posterior tailbud region. However, the specific biological determinant that integrates these cellular processes during vertebrate tail bud elongation has remained unclear. Here, we demonstrate that the spatiotemporal heterogeneity of chromatin condensation states governs both cell fate and cell motility within the mesodermal progenitor zone (MPZ) and the presomitic mesoderm (PSM) during zebrafish tail bud elongation. By integrating biological experiments with computational modeling, we reveal that nuclear volume heterogeneity, mediated by chromatin remodeling, acts as an active driver of collective cell migration. Reducing chromatin condensation heterogeneity in the MPZ through modulation of chromatin modifications impairs collective cell movements and delays tailbud elongation. Single cell ATAC-seq and RNA-seq analyses reveal that global chromatin accessibility heterogeneity and region-specific chromatin remodeling differences in the PSM and MPZ are functionally associated with gene expression pathways involving in cell lineage specification. These findings establish a molecular connection between cell fate determination and tissue-scale cellular dynamics. Collectively, our findings unveil a unifying framework for understanding the coordination of cell fate determination and large-scale cellular movements during tissue morphogenesis.

## INTRODUCTION

The effective elongation of the vertebrate body axis depends on the precise coordination of cell movements within the posterior tailbud region, ensuring the orderly formation of tissues^1–3^. In zebrafish (*Danio rerio*), disruptions to these tailbud cell movements lead to delayed or impaired axis elongation, highlighting the critical role of collective cell dynamics in embryonic development^4–6^. While many genetic and chemical gradients—including Fgf, Wnt, and Notch signaling pathways—have been implicated in tailbud morphogenesis^7–9^, the mechanisms by which genetic or cell fate-determining molecules integrate with collective cell motion to drive large-scale morphogenetic processes remain poorly understood.

Among the diverse subcellular regulators, the nucleus emerges as a central player due to its substantial size, mechanical rigidity, and critical role in gene expression regulation^10–13^. Beyond its well-established function in orchestrating transcriptional programs, the nucleus directly influences cellular mechanics, thereby affecting collective tissue behaviors^14,15^. Notably, chromatin regulation— encompassing processes such as chromatin remodeling, histone modifications, and higher-order organization—not only shapes nuclear architecture but also fine-tunes the mechanical properties of the nucleus^16–18^. Furthermore, changes in nuclear lamina composition can either reinforce or relax the nuclear envelope, directly impacting how cells perceive and transmit mechanical forces^19–22^. Elucidating how these chromatin- and lamina-mediated modifications reconfigure nuclear properties, and consequently drive tissue-level dynamics, is crucial for a comprehensive understanding of morphogenic processes underlying both embryonic development or tumor metastasis.

From a biophysical perspective, cells within dense tissues can transition between solid-like or fluid-like states under varying conditions, a phenomenon known as jamming or unjamming transitions^23^. During zebrafish tailbud development, continuous posterior elongation depends on active cell migration and precise alteration of jamming transitions from the mesodermal progenitor zone (MPZ) and the presomitic mesoderm (PSM)^4,5^. While traditional models emphasize cell density, cell–cell adhesion, and cell shape as primary determinants of jamming transitions^13^, the specific biological mechanisms that govern cellular movement within tissues—and link cell fate determination to tissue morphogenesis— remain poorly understood.

In this study, we investigated how chromatin remodeling- and lamina-mediated regulation of nuclear architecture coordinate cellular dynamics and cell fate decision during zebrafish body axis elongation. Through live imaging combined with pharmacological inhibition of chromatin remodeling, we demonstrate that spatiotemporal nuclear volume heterogeneity is tightly correlated with cell motility in MPZ and PMS. Restricting nuclear volume heterogeneity by modulating chromatin modifications impairs collective cell rearrangements and delays axis elongation, establishing a direct link between subcellular modifications and tissue-scale mechanical outcomes. Simplified computational modeling reveals that nuclear volume fluctuations, shaped by chromatin remodeling, function as an active driver for tissue fluidity. Single-cell epigenetic profiling further demonstrates that global and region-specific differences in chromatin remodeling modulate key pathways governing cellular motility and cell lineage specification, providing a mechanistic connection between cell fate regulation and large-scale cell dynamics. Together, these results underscore the central role of chromatin remodeling in tailbud morphogenesis and offer a fresh perspective on how subcellular modifications orchestrate vertebrate morphogenesis.

## RESULTS

### Spatiotemporal heterogeneity of chromatin condensation state decreases during zebrafish body axis elongation

To explore the role of nuclear dynamics in collective cell movement during zebrafish tailbud development, we analyzed cell nuclei in zebrafish embryos at the 10-somite stage using DAPI staining. Quantitative analysis revealed significant differences in nuclear volume between cells in MPZ and PSM. Specifically, the average nuclear volume was significantly higher in MPZ cells compared to PSM cells. Notably, MPZ cells displayed broader nuclear volume distribution, indicating greater dynamism compared to PSM cells (Fig. 1a and b). Assessment of nuclear volume variability, quantified by standard deviation, confirmed a reduction in nuclear volume heterogeneity during developmental transition from MPZ to PSM (Fig. 1c). These findings demonstrate that the spatial heterogeneity in nuclear volume progressively diminishes during zebrafish body axis elongation.

**Figure 1.**
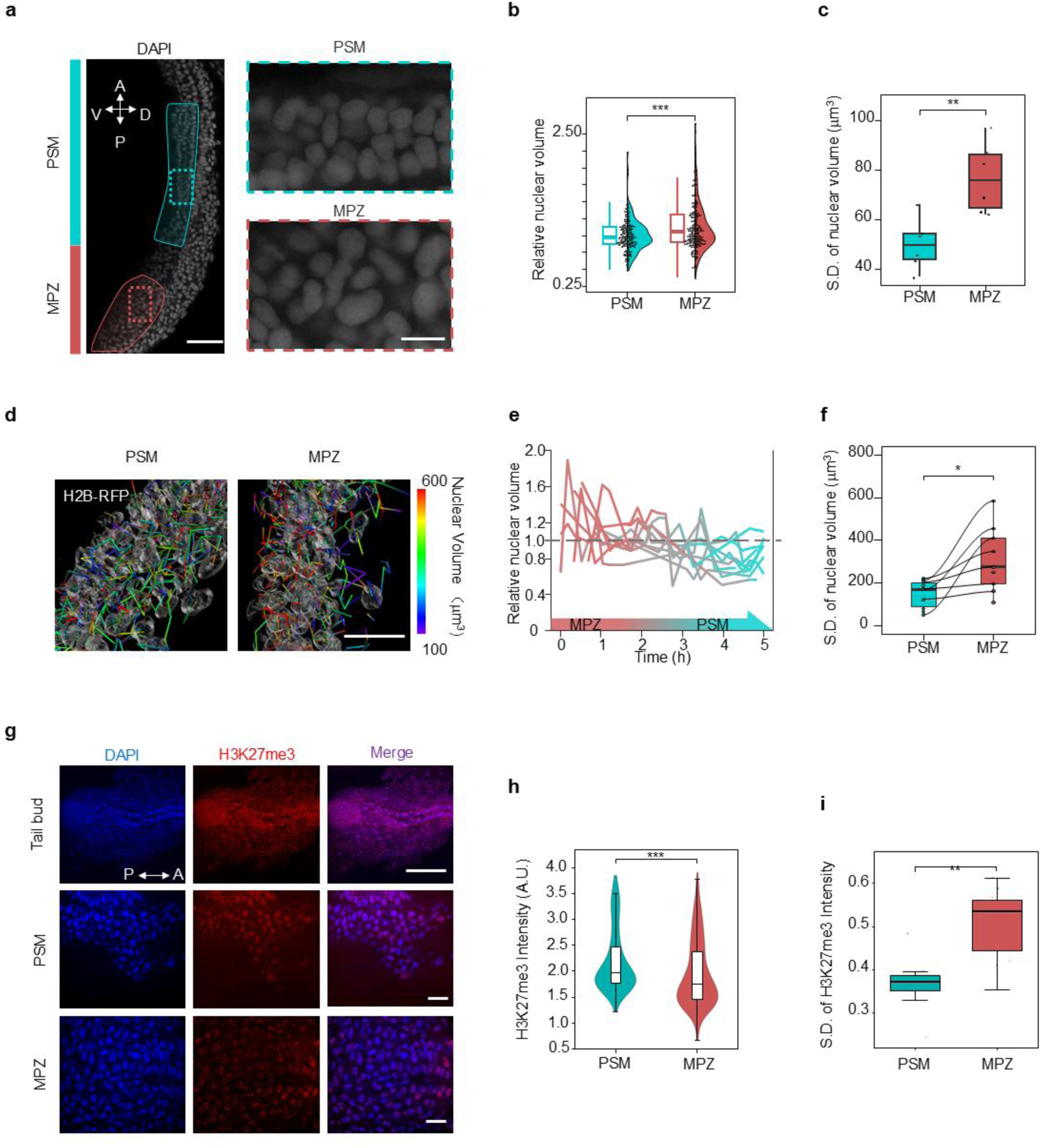
Nuclear volume fluctuations during zebrafish embryo body axis elongation. (a) Whole-embryo DAPI staining in the sagittal plane of the zebrafish tailbud, showing the mesodermal progenitor zone (MPZ, outlined in red) and the presomitic mesoderm (PSM, outlined in cyan). Enlarged insets display nuclear organization within the MPZ and PSM regions. Scale bar: 50 µm, zoom: 10µm. (b) Quantification of nuclear volumes in the MPZ and PSM regions derived from data in (a), indicating that nuclear volumes are significantly larger in MPZ cells compared to PSM cells (n=209(PSM), 221(MPZ), ***p < 0.001, t test). (c) Statistical analysis of nuclear volume standard deviation derived from data in (a), showing greater variability in nuclear volume in MPZ cells than in PSM cells (n=6(PSM), 6(MPZ), **p < 0.01, t test). (d) Live-cell imaging of zebrafish embryos injected with H2B-RFP mRNA, showing labeled nuclei in both MPZ and PSM regions. Scale bar: 50 µm. (e) Relative nuclear volume changes of 10 individual cells tracked as they migrate from the MPZ to the PSM, illustrating a reduction in nuclear volume fluctuations during this transition. (f) Standard deviation of nuclear volume for the tracked cells in (e), showing a significant decrease in nuclear volume variability as cells move from MPZ to PSM (n=10(PSM), 9(MPZ), *p < 0.05, t test). (g) Whole-embryo DAPI (blue) and H3K27me3 (red) staining in wide type zebrafish embryo, showing the mesodermal progenitor zone (MPZ) and presomitic mesoderm (PSM) regions. This staining was used for calculating nuclear volume in the subsequent panels. Scale bar: 50 µm, zoom: 25µm. (h) Quantification of H3K27me3 fluorescence intensity in the MPZ and PSM regions of g. (**p < 0.01, t-test). (i) Quantification of H3K27me3 fluorescence intensity in the MPZ and PSM regions of g (**p < 0.01, t-test).

To further investigate the dynamic changes in nuclear volume during the transition of individual cells from MPZ to PSM, we performed live imaging of H2B-RFP-labeled zebrafish embryos to monitor nuclear volume changes in tail bud cells across the 8-13 somite stages. (Fig. 1d, Supplemental Movie S1). Time-lapse imaging analysis revealed a progressive reduction in both the average value of nuclear volume and nuclear volume fluctuations as cells migrated from MPZ to PSM (Fig. 1e). Quantitative assessment of nuclear volume variability, as measured through standard deviation analysis, further confirmed a significant decrease in nuclear volume variance during this developmental transition (Fig. 1f). Together, these results provide evidence that the spatiotemporal heterogeneity of nuclear volume progressively decreases in parallel with the advancing development of the zebrafish body axis.

Given the critical role of chromatin condensation state in regulating nuclear mechanics and influencing nuclear volume, we investigated the epigenetic landscape during zebrafish tail bud development. Immunofluorescence staining using an antibody against H3K27me3, a marker of repressive chromatin, revealed significantly higher levels of H3K27me3 in MPZ cells compared to PSM cells (Fig. 1g and h). Furthermore, the distribution of H3K27me3 signal intensity in each cells exhibited distinct patterns between these MPZ and PMS. Quantitative analyses of standard deviation and coefficient of variation demonstrated that H3K27me3 signal intensity was more uniform in the PSM cells, with lower variance. In contrast, MPZ cells displayed greater variability in H3K27me3 signal intensity (Fig. 1i). These results demonstrate that the heterogeneity of chromatin condensation state is reduced during the development of zebrafish tail bud.

### Spatiotemporal heterogeneity of chromatin condensation state decreases in jammed tissues

Previous studies have reported that during the development from MPZ to PSM, the collective movement of cells decreases, a process known as jamming transitions, which plays a critical role in zebrafish body axis elongation^5^. To explore the relationship between chromatin condensation and jamming transitions, we utilized MDCK cell sheets—a well-established model for studying tissue fluidity transition (Fig. 2a). By employing GFP-NLS–labeled MDCK cells, we combined live-cell imaging with Particle Image Velocimetry (PIV) to quantify cell motility in jammed (low motility) and unjammed (high motility) regions, while simultaneously comparing nuclear volume distributions between these states (Fig. 2b–d, Supplemental Movie S2). Our analyses demonstrated that the average nuclear volume in jammed regions was significantly smaller than that in unjammed regions. Furthermore, measures of dispersion—including standard deviation and coefficient of variation— revealed that nuclear volumes of individual cells in the jammed states were markedly more uniform compared to those in unjammed states (Fig. 2e–f). These findings highlight a clear distinction in nuclear volume heterogeneity between jammed and unjammed tissues, suggesting a potential link between tissue fluidity and nuclear volume.

**Figure 2.**
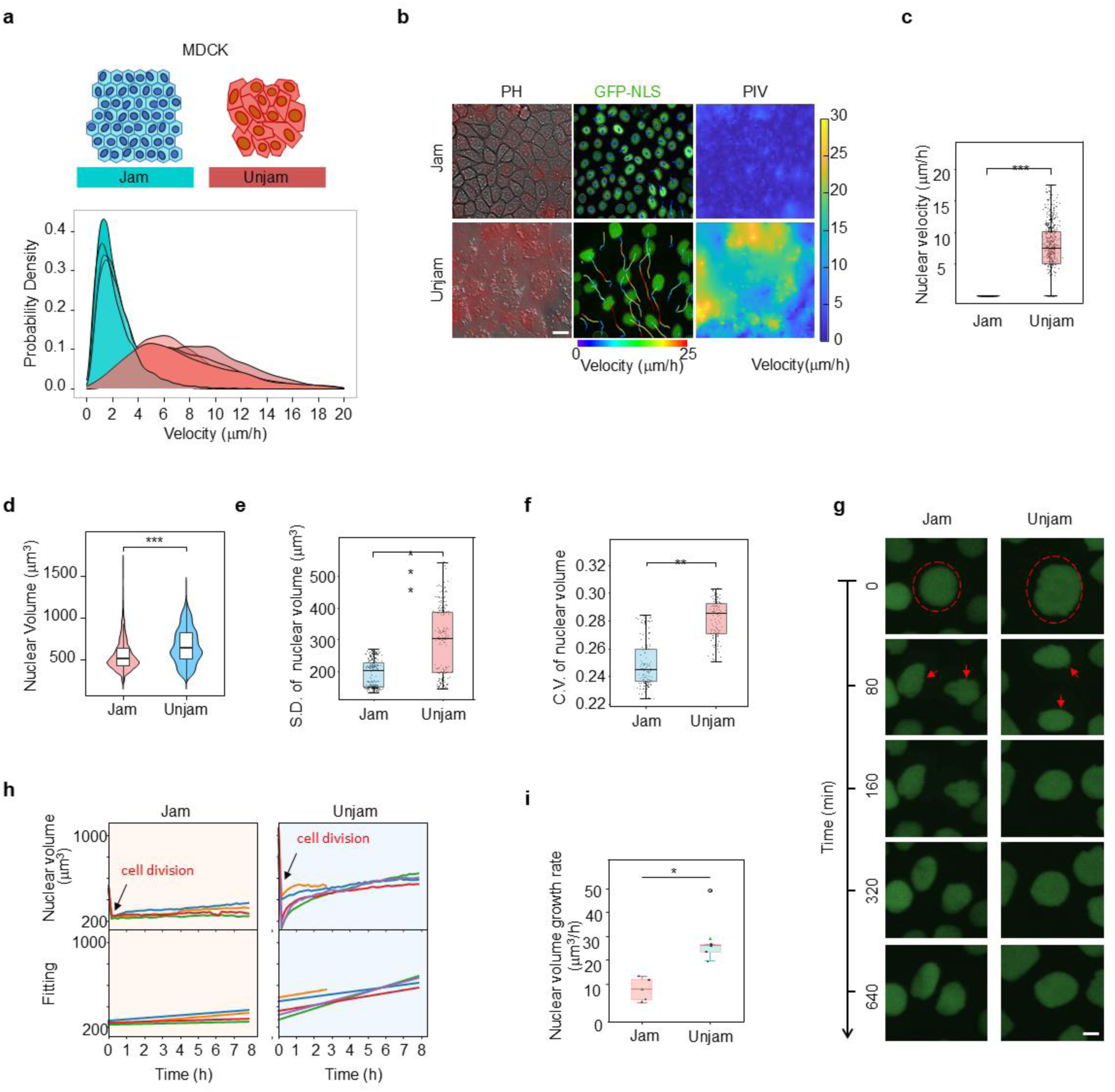
Nuclear volume fluctuations during jamming transitions in epithelial tissues. (a) Schematic of Madin-Darby canine kidney (MDCK) cell monolayers displaying two distinct states: jammed (blue) and unjammed (red). Below: Probability density of cell velocities in the MDCK monolayer, demonstrating that MDCK cells exhibit both jammed and unjammed states, making them a suitable model for studying jamming transitions. (b) Live-cell imaging of MDCK-GFP-NLS cells showing nuclear morphology in both jammed and unjammed states (left) and particle image velocimetry (PIV) analysis (right). Scale bar: 20 µm. (c) nuclear velocities in jammed and unjammed regions, showing that nuclear velocities in unjammed regions are larger than jamming velocities (***p < 0.001, t test). (d) Distribution of nuclear volumes in jammed and unjammed regions, showing that nuclear volumes are not only larger in the unjammed region but also exhibit a broader distribution range (n=19043(jammed), 11787(unjammed), ***p < 0.001, t test). (e) Comparison of nuclear volume standard deviation between jammed and unjammed regions (n=312(jammed), 184(unjammed), ***p < 0.001, t test). (f) Comparison of nuclear volume coefficient of variation between jammed and unjammed regions (n=312(jammed), 184(unjammed), ***p < 0.001, t test). (g) Live-cell imaging of post-division nuclear volume over time in the stuck and unstuck states demonstrates the dynamics of nuclear volume growth. (h) Statistics and growth rate fitting for tracking the volume of single cells after cell division. (i) Quantitative statistics of cell growth rates after cell division in jammed regions and unjammed regions of (h).

To investigate the underlying mechanisms driving the observed differences in nuclear volume between jammed and unjammed states, we monitored nuclear volume dynamics in individual cells throughout the cell cycle (Figure 2g). Our data revealed that cells in the jammed regions exhibit a significantly slower rate of nuclear volume growth starting from the post-mitotic stage compared to those in the unjammed regions (Fig. 2h–i). Moreover, the growth rate in unjammed regions displayed considerably greater heterogeneous than in jammed regions. (Fig. 2i). This disparity in nuclear volume growth dynamics appears to be the key factor contributing to the more stable and uniform nuclear volume distribution observed in jammed tissues. Collectively, these findings suggest that variations in nuclear volume growth dynamics following cell division are attenuated in jammed tissues.

We next investigated the chromatin condensation state in jammed and unjammed regions of MDCK cell sheets. Immunofluorescence staining using an antibody against H3K9me3, a heterochromatin marker, revealed significantly higher levels of H3K9me3 in unjammed regions compared to jammed regions (Fig. 3a-b). Furthermore, the distribution of H3K9me3 signal intensity in individual cells exhibited distinct patterns between these regions. Quantitative analyses of standard deviation and coefficient of variation demonstrated that H3K9me3 signal intensity was more uniform, with lower variance, among jammed cells. In contrast, unjammed regions displayed greater variability in H3K9me3 signal intensity (Fig. 3c). Similar trends were observed for another heterochromatin marker, HP1-α, where the variability in signal intensity was significantly higher in unjammed regions compared to jammed ones (Fig. 3d-f). Collectively, these results suggest that fluctuations in chromatin condensation state are attenuated in jammed tissues.

**Figure 3.**
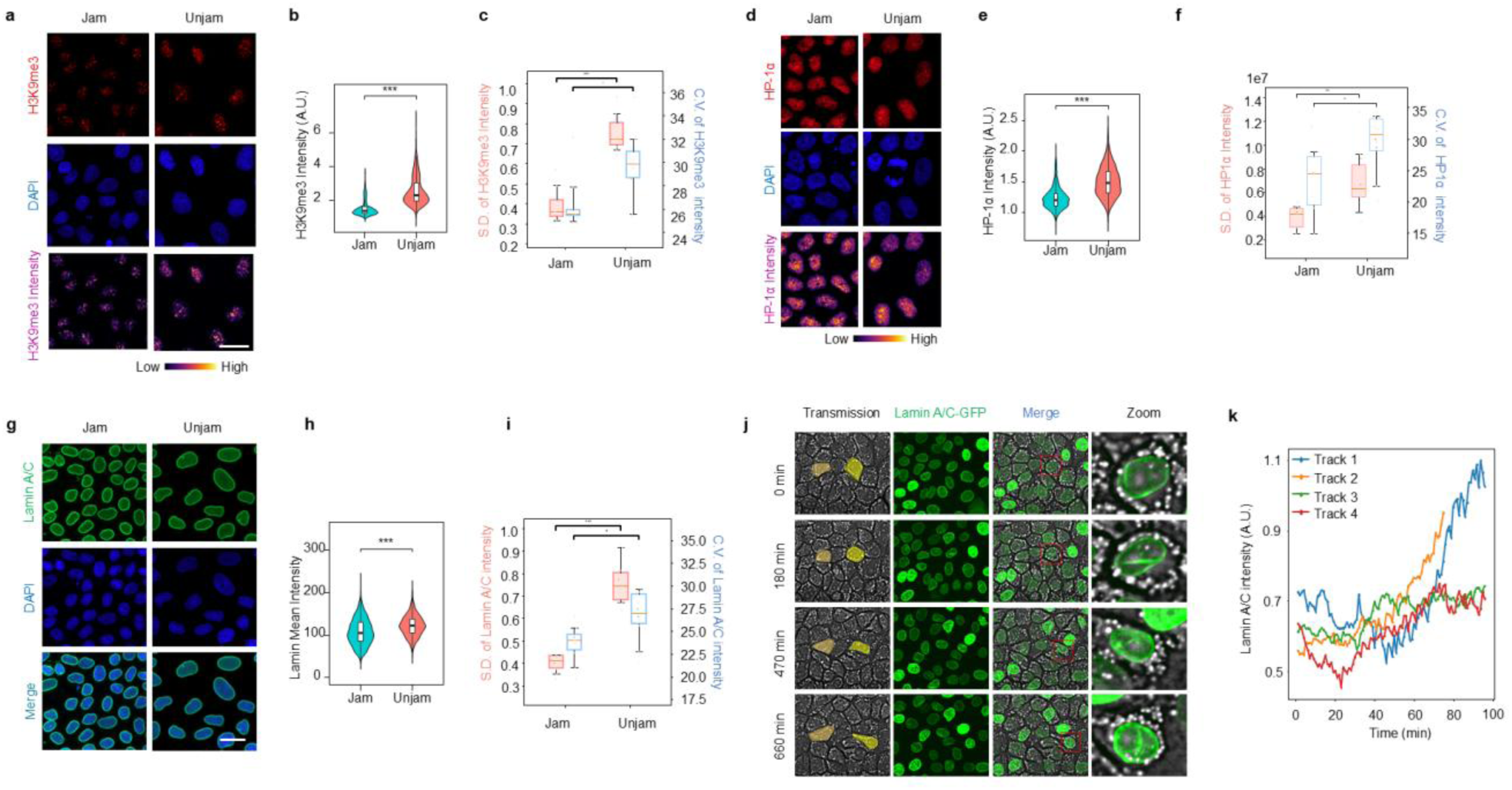
Chromatin condensation and nuclear lamina fluctuations during jamming transitions. (a) Immunofluorescence staining of H3K9me3 (red) and DAPI (blue) in jammed and unjammed regions, indicating levels of chromatin condensation. Color bar represents fluorescence intensity from low (left) to high (right). Scale bar: 20 µm. (b) Quantification of H3K9me3 fluorescence intensity in jammed and unjammed regions, showing significantly higher fluorescence intensity in the unjammed regions compared to the jammed regions. (n=10(jammed), 10(unjammed), **p < 0.01, t-test). (c) Quantification of H3K9me3 fluorescence intensity standard deviation and coefficient of variation (CV) in jammed and unjammed regions, showing higher variance and lower kurtosis in the jammed state compared to the unjammed state (n= 10(jammed), 10(unjammed), **p < 0.01, ***p < 0.001, t test). (d) Immunofluorescence staining of HP1-α (red) and DAPI (blue) in jammed and unjammed regions, displaying the distribution of HP1-α under different states. Color bar represents fluorescence intensity. Scale bar: 20 µm. (e) Quantification of HP1-α fluorescence intensity in jammed and unjammed regions, showing significantly higher fluorescence intensity in unjammed regions compared to jammed regions (n=10(jammed), 12(unjammed), **p < 0.01, t-test). (n=10(jammed), 12(unjammed), **p < 0.01, t-test). (f) Quantification of HP1-α fluorescence intensity standard deviation and coefficient of variation (CV), showing that in the jammed state, HP1-α exhibits significantly higher intensity variance and lower kurtosis compared to the unjammed state (n= 10(jammed), 12(unjammed), ***p < 0.001, t test). (g) Immunofluorescence staining of Lamin A/C (green) and DAPI (blue) in jammed and unjammed regions. Scale bar: 20 µm. (h) Quantification of Lamin A/C fluorescence intensity, showing significantly higher fluorescence intensity in unjammed regions compared to jammed regions (n=7(jammed), 6(unjammed), **p < 0.05, t-test). (i) Comparison of Lamin A/C fluorescence intensity standard deviation and coefficient of variation (CV) between jammed and unjammed states (n= 7(jammed), 6(unjammed), **p < 0.01.) (j) Time-lapse images of live MDCK cells expressing Lamin A/C-GFP, showing dynamic changes in Lamin A/C during cell intercalation. Increased Lamin A/C expression is observed in response to intense nuclear deformation during intercalation events. Scale bar: 20 µm, Zoom: 10 µm. (k) Quantification of Lamin A/C fluorescence changes in single-cell tracking of the occurrence of the T1 transition (n=4).

Nuclear lamina is a key structural component involved in regulating chromatin condensation state. We next investigated the dynamics of the nuclear lamina between jammed and unjammed states. Immunofluorescence analysis using an antibody against Lamin A/C, a principal constituent of the nuclear lamina, revealed that the average nuclear lamina intensity was significantly higher in unjammed regions compared to jammed regions (Fig. 3g and h). Furthermore, the variance and coefficient of variation of nuclear lamina signals in individual cells were greater in unjammed cells compared to jammed cells (Fig. 3i). To further explore nuclear lamina dynamics, we employed MDCK cells expressing Lamin A/C-GFP and monitored nuclear lamina intensity in jammed and unjammed regions. Live-cell imaging demonstrated that changes in nuclear lamina signal intensity predominantly occurred in regions undergoing T1 transitions, where nuclei undergo large deformation to facilitate tissue fluidization (Fig. 3j and k, Supplemental Movie S2). Collectively, these findings demonstrate a tight correlation between spatiotemporal fluctuations in chromatin condensation and tissue fluidity states.

### Chromatin remodeling dictates tissue fluidity

Consequently, we sought to determine the role of chromatin condensation states in regulating tissue fluidity. To manipulate chromatin modifications, we employed the histone deacetylase inhibitor Trichostatin A (TSA) and the histone acetyltransferase inhibitor Garcinol (Gar) in MDCK cell sheets. Immunofluorescence staining revealed that TSA treatment led to a reduction in HP1-α intensity, whereas Garcinol increased it (Fig. 4a-c). Notably, both treatments significantly reduced the spatial heterogeneity of HP1-α intensity, as indicated by decreased standard deviation and coefficient of variation (Fig. 4b-d). Further analysis of nuclear volume demonstrated that TSA treatment significantly increased nuclear volume, while Garcinol treatment reduced it; importantly, both treatments lowered the standard deviation of nuclear volume, indicating a decrease in spatial heterogeneity (Fig. 4e and f).

**Figure 4.**
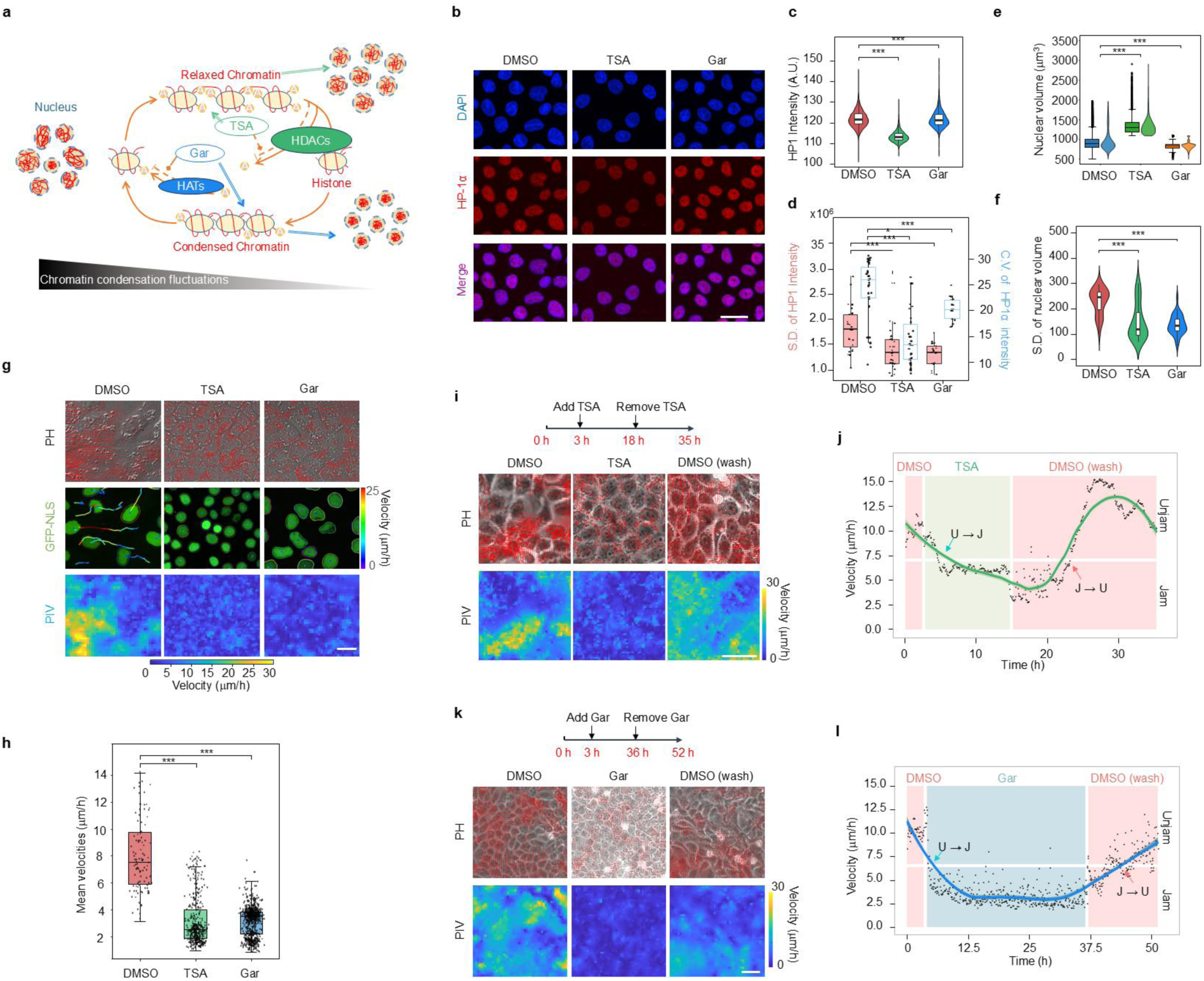
Jamming transitions require nuclear volume fluctuations. (a) Schematic illustrating the effects of TSA and Garcinol on chromatin condensation. By modulating chromatin structure, these drugs ultimately aim to control nuclear volume fluctuations: TSA (a histone deacetylase inhibitor) relaxes chromatin by inhibiting HDAC, increasing nuclear volume stability, while Garcinol (a histone acetyltransferase inhibitor) promotes chromatin condensation by inhibiting HAT, reducing nuclear volume fluctuations. (b) Immunofluorescence staining of HP1-α (red) and DAPI (blue) at DMSO, TSA (20uM) and Garcinol(10uM). Scale bar: 20 µm. (c) Quantification of HP1-α fluorescence intensity (n=9705(DMSO), 4091(TSA 20nM), 12468(Garcinol 10uM), ***p < 0.001, t-test). (d) HP1-α fluorescence intensity standard deviation and coefficient of variation (CV) of (b) (n=29(DMSO), 30(TSA 20nM), 39(Garcinol 10uM), ***p < 0.001, t-test). Drug-washout experiment for TSA. (e) Quantification of nucleus volume of (b) (n=9705(DMSO), 4091(TSA 20nM), 12468(Garcinol 10uM), ***p < 0.001, t-test). (f) Quantification of nucleus volume standard deviation (n=29(DMSO), 30(TSA 20nM), 39(Garcinol 10uM), ***p < 0.001, t-test). (g) Live-cell images of MDCK-GFP-NLS cells after treatment with DMSO (control), TSA, and Garcinol, followed by PIV analysis to assess cell motility. Scale bar: 20 µm. (h) Quantitative statistics of nuclear velocities in DMSO (control), TSA and Garcinol of (g). (i, k) Time-lapse images of MDCK cells showing the reversible effects of TSA (i) and Garcinol (k) on cell motility and nuclear volume fluctuations. The upper timeline indicates the timing of drug addition and removal, with corresponding live-cell images and PIV analyses displayed. (j, l) Cell velocity profiles over time during and after TSA (i) and Garcinol (k) treatment, showing a significant decrease in cell motility during drug treatment, with a gradual recovery in motility after drug removal.

We further investigated the effects of chromatin condensation state heterogeneity on tissue fluidity in cells treated with TSA and Garcinol. Live-cell imaging combined with PIV analysis demonstrated that both TSA and Garcinol treatments significantly reduced cell motility (Fig. 4g and h). Long-term time-lapse imaging revealed that unjammed cells treated with either TSA or Garcinol gradually transitioned to a jammed-like state, characterized by reduced motility. Importantly, upon drug washout, cell motility returned to pre-treatment levels, highlighting the reversibility of chromatin modification– mediated regulation of tissue fluidity (Fig. 4i–l). Analysis of cell density during the imaging period confirmed that the observed changes in motility were not attributable to fluctuations in cell density (Supplementary Fig. 4). Collectively, these results establish that chromatin remodeling plays a pivotal role in determining tissue fluidity.

### Chromatin remodeling-mediated nuclear volume fluctuations act as a self-enhancing driver for cellular jamming transitions

To further validate our experimental findings and elucidate the mechanistic role of the chromatin remodeling in tissue fluidity, we developed two theoretical models that integrate subcellular nuclear dynamics with classical jamming transitions. Jamming transitions are typically modeled using assemblies of 2D disks or 3D spheres for passive particles^24,25^, or an assembly of irregular polygons with self-propelled forces for active cells^23,26^. To better understand the role of chromatin remodeling-mediated nuclear volume fluctuations in jamming transitions, we propose two theoretical models: (i) In the first model (the effective disk model, EDM), nuclei are considered as 2D disks with an effective size λ_*i*_. Without loss of generality, their complex interactions are modeled using a modified long-range Lennard-Jones (LJ) potential, and their motion follows Newtonian dynamics. Nuclear volume fluctuations are incorporated through a distribution of effective disk sizes with a standard deviation 𝛿λ. (ii) In the second model (the self-propelled Voronoi model, SPVM), cells are described as tightly packed irregular polygons generated by Voronoi tessellation, which move following over-damped Langevin dynamics with a self-propelled active force. Previous studies have established that the behavior of the SPVM is governed by a shape index *p* that characterizes the shape of cells^23^. Here, we assume that the nuclear volume fluctuations correspond to a distribution of preferred cell shapes with a mean value *p* and a standard deviation 𝛿𝑝, which are mechanically coupled through cytoskeletal filaments and additional pathways, including focal adhesions and nuclear lamina interactions^27^. This coupling is supported by the experimentally observed correlation between the shape index and nuclear volume (Supplementary Fig. xx). Further details of the models are provided in Materials and Methods. In both models, the jammed and unjammed states are determined by a dynamical order parameter, the diffusivity (Fig. 5a and b). The jamming phase diagram is universally controlled by two parameters: in the EDM, these are the size fluctuation 𝛿λ and the area fraction 𝜙 (Fig. 5c); in the SPVM, they are the shape fluctuation 𝛿𝑝 and the mean shape index *p* (Fig. 5d). In both cases, nuclear volume fluctuations, represented by 𝛿λ and 𝛿𝑝 respectively, can induce fluidity transitions when the other parameter is fixed, consistent with our experimental observations. To further explore the role of nuclear volume fluctuations as an active source, we perform additional simulations for the EDM, allowing disk sizes to fluctuate temporally during dynamics. This “breathing-like” motion introduces additional degrees of freedom^28^, further fluidizing the system with an increased diffusivity (Fig. 5f). These numerical analyses confirm that chromatin remodeling-mediated nuclear volume fluctuations serve as a self-enhancing driver for cellular jamming transitions.

**Figure 5.**
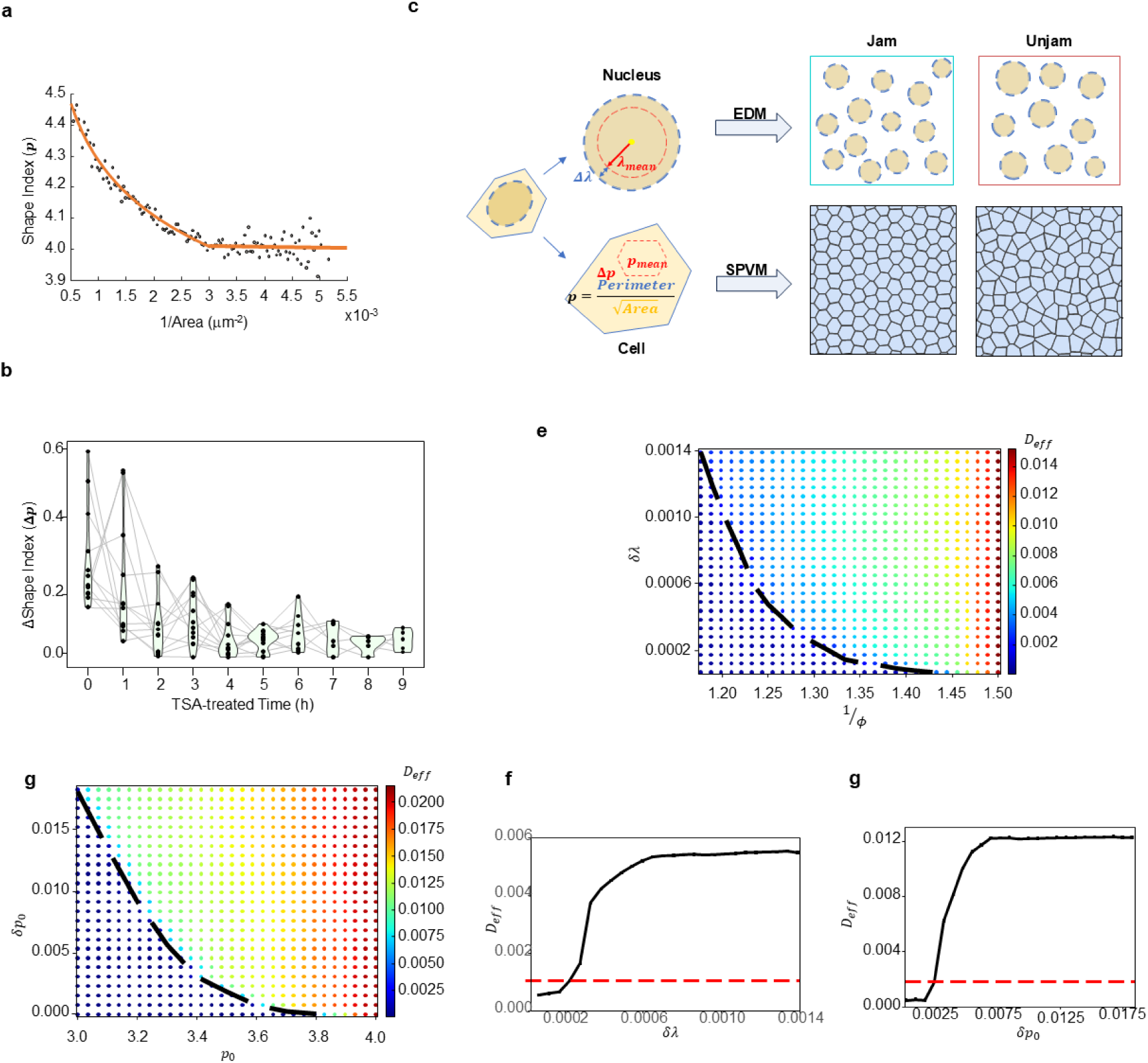
Numerical modeling of the tissue fluidity transition, by the (a-d) EDM and (e-h) SPVM. (a) Diffusivity D as a function of the size fluctuation 𝛿λ, with the area fraction 𝜙=?? fixed. The dashed red line is the threshold D=0.001 as the jamming criterion. The blue line corresponds to the data obtained with “breathing-like” motion. (e) Diffusivity D as a function of the shape fluctuation 𝛿𝑝, with the mean shape index p=?? fixed. (b, f) The phase diagrams and typical (c, g) unjammed and (d, h) jammed configurations are presented for both models.

### Chromatin remodeling integrates collective cell movement and cell fate determination during zebrafish body axis elongation

During zebrafish tail bud development, the transition of cell fate is orchestrated by the antagonistic interplay between *sox2*, which maintains pluripotency in the MPZ, and *tbxta*, which drives commitment to the PSM. The dynamic gene expression gradients of *sox2* and *tbxta* establish a bistable switch that precisely coordinates the MPZ-to-PSM transition^29,30^. To explore the role of chromatin remodeling on this cell fate decision process, we first examined the epigenetic landscape by integrating published single-cell ATAC-seq (scATAC-seq) and single-cell RNA-seq (scRNA-seq) data from developing zebrafish tail bud. Our genome-wide comparison of all detected chromatin accessibility peaks revealed that MPZ cells exhibit significantly higher variance in chromatin accessibility compared to PSM cells (Fig. 6a-b), indicating a more heterogeneous epigenetic state within the MPZ population. This finding suggests that epigenetic variability may play a critical role in regulating cell fate plasticity during early developmental stages.

**Figure 6.**
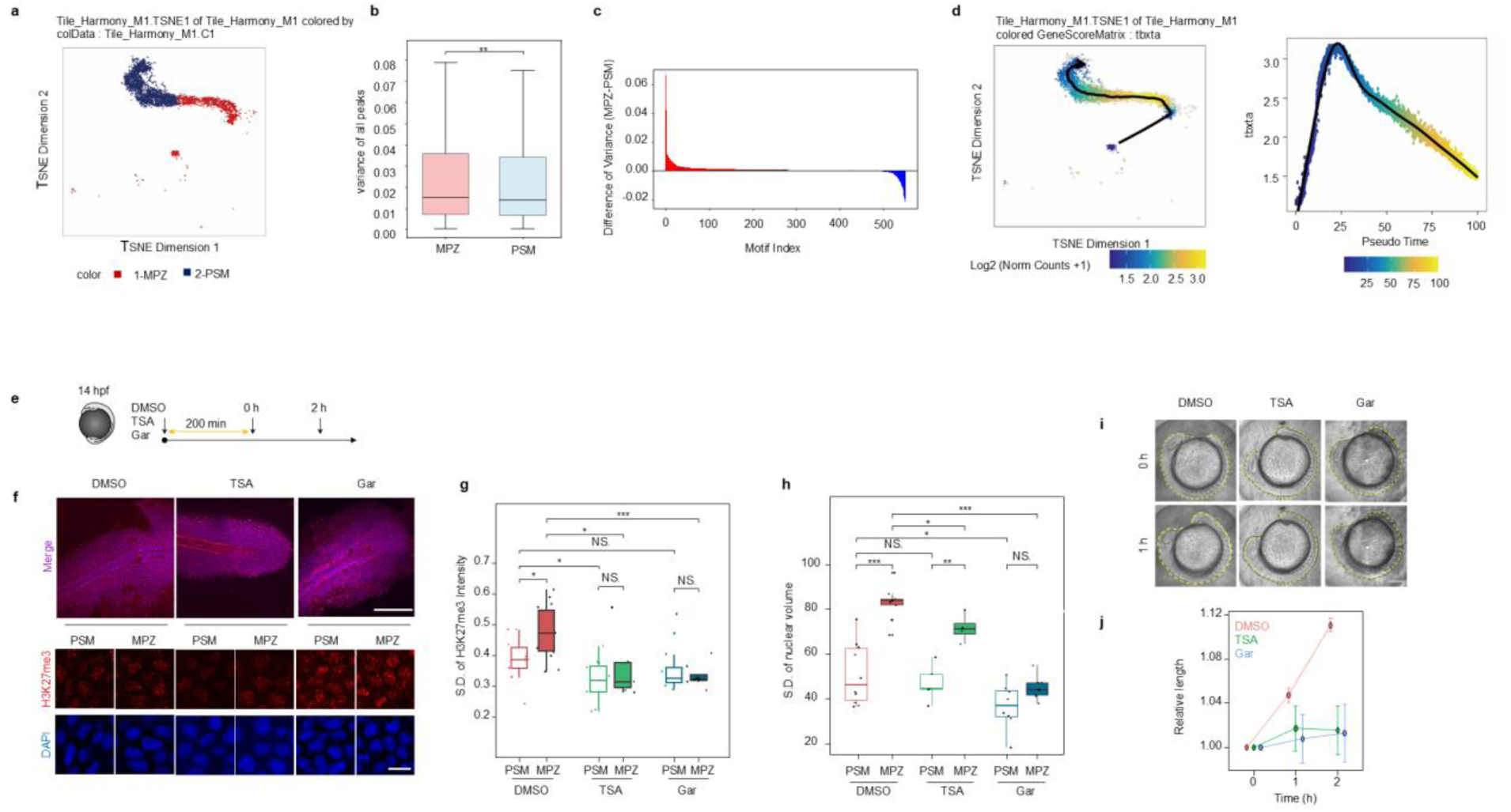
Nuclear volume fluctuations facilitate zebrafish body axis development. (a)A t-SNE projection of single-cell ATAC-seq data from 14-hpf zebrafish embryos, highlighting two distinct subpopulations: mesodermal progenitor zone (MPZ, red) and presomitic mesoderm (PSM, blue). This separation reflects region-specific differences in chromatin accessibility underlying progenitor versus somitogenic identities. (b) Quantitative statistics of variance in scATAC-seq peak accessibility across all detected peaks in the mesodermal progenitor zone (MPZ) and presomitic mesoderm (PSM). (c) The bar plot shows the difference in motif-level scATAC-seq variance between the MPZ and PSM regions in zebrafish, red bars indicate motifs whose variance is higher in MPZ than in PSM, whereas blue bars represent motifs whose variance is higher in PSM. (d) Additional scATAC-seq–based dimensionality reduction plots and density curves demonstrating progressive changes in chromatin accessibility of tbxta along the pseudotime axis. (e)Schematic illustration showing the drug treatment protocol, including the time points of TSA and Garcinol addition and removal. (f) Whole-embryo DAPI (blue) and H3K27me3 (red) staining in zebrafish embryos following TSA and Garcinol treatment, showing the mesodermal progenitor zone (MPZ) and presomitic mesoderm (PSM) regions. This staining was used for calculating nuclear volume in the subsequent panels. n=103(DMSO PSM), 88(DMSO MPZ), 184(TSA PSM), 177(TSA MPZ), 140(Garcinol PSM), 181(Garcinol MPZ) Scale bar: 50 µm, zoom: 10µm. (g) Quantification of nuclear volume variance in the MPZ and PSM regions post-treatment. TSA and Garcinol treatment significantly reduced the variance in nuclear volume in both regions, with a more pronounced reduction in the MPZ (n=5 (TSA), 7 (Garcinol), 10(DMSO), **p < 0.01, t-test). (h) Quantification of H3K9me3 fluorescence intensity variance in the MPZ and PSM regions post-treatment. TSA and Garcinol treatment significantly reduced the variance in nuclear volume in both regions, with a more pronounced reduction in the MPZ (n=17 (TSA), 20(Garcinol), 25(DMSO), **p < 0.01, t-test). (i) Live-cell imaging of zebrafish embryos treated with TSA and Garcinol, showing developmental changes in the body axis at different time points. Scale bar: 200 μm. (j) Quantification of relative body axis length over time, showing that TSA and Garcinol treatment significantly inhibits body axis elongation, with a reduced elongation rate compared to the control group (n=3, **p < 0.01, t-test).

During zebrafish tail bud development, *sox2* and *tbxta* are two genes involved in the maintenance of progenitor states and driving early mesoderm specification, while *tbx16* and *zeb1* contribute to somitogenesis and the regulation of mesenchymal cell phenotypes^31–33^. Detailed analyses of these key regulatory loci revealed distinct chromatin accessibility patterns: *sox2* and *tbxta* exhibited enhanced accessibility in the MPZ, whereas *tbx16* and *zeb1* showed increased accessibility in the PSM (Supple mental Fig 7). These chromatin accessibility profiles align with our observations of cellular motion transitions from MPZ to PSM, where MPZ cells displayed greater nuclear volume fluctuations and fluid-like motility, contrasting with the PSM cells which exhibited reduced nuclear volume fluctuations and solid-like motility. These findings suggest a mechanistic connection between chromatin accessibility, cell fate decisions, and cell movement during zebrafish tail bud development.

**Fig 7.**
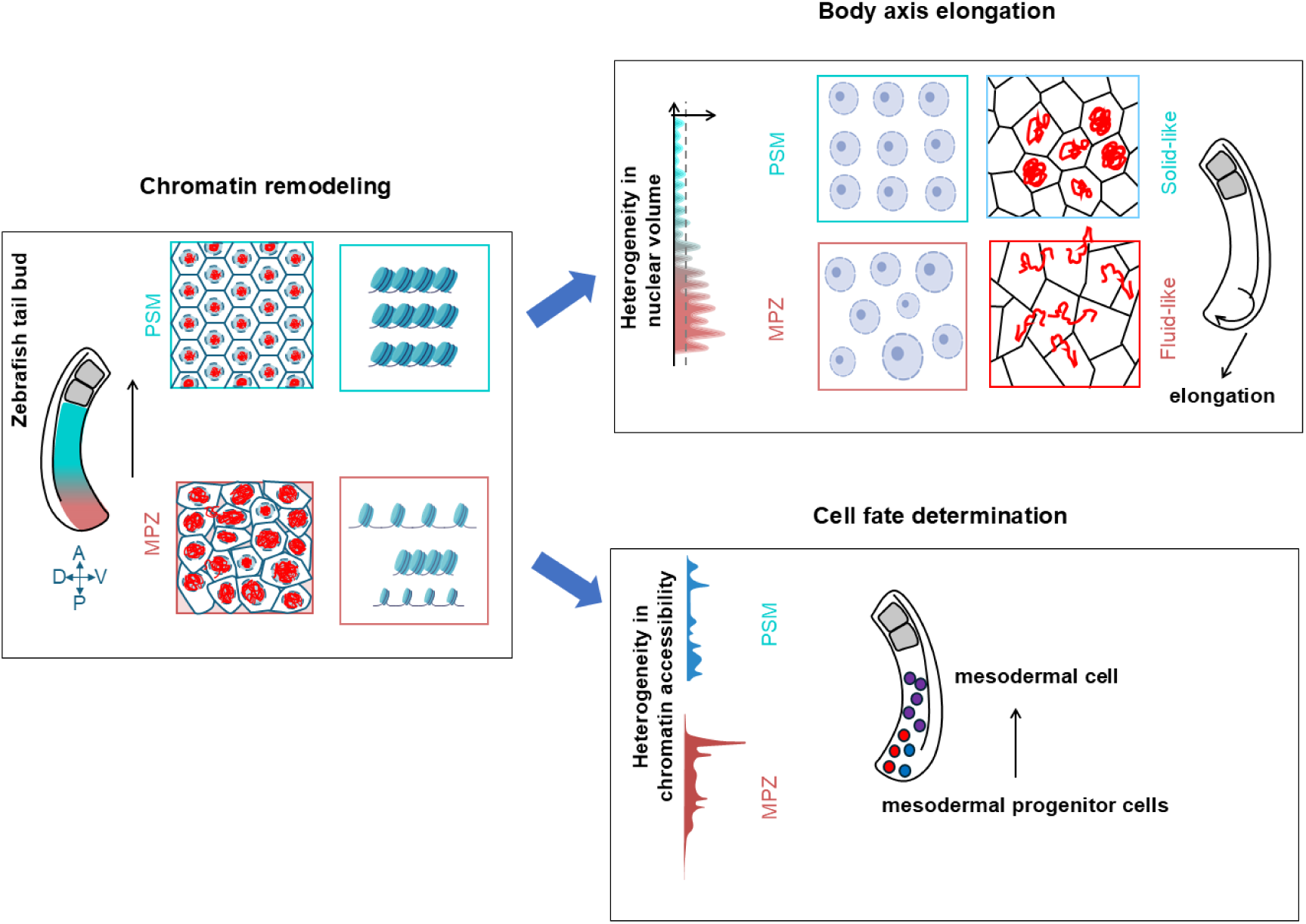
Chromatin remodeling drive body axis elongation and cell fate transitions. Schematic summarizing the interplay between the mesodermal progenitor zone (MPZ) and the presomitic mesoderm (PSM) during zebrafish tail axis extension. In the MPZ, more accessible chromatin and greater nuclear volume heterogeneity support a fluid-like, unjammed state, whereas the PSM adopts a jammed, solid-like configuration linked to tighter chromatin organization. This chromatin-driven remodeling influences both tissue fluidity and nuclear architecture, thereby affecting the rate and efficiency of body axis elongation. Single-cell analyses further reveal distinct chromatin accessibility landscapes for MPZ and PSM, indicating that epigenetic dynamics not only govern cell motility but also coordinate cell fate decisions throughout tailbud development.

Building upon this epigenetic groundwork, we modulated chromatin modification states by treating the tailbud region of zebrafish embryos at 14 hpf with TSA and Gar (Fig. 6c-d). Through quantitative analysis of H3K27me3 fluorescence intensity and nuclear volume, we observed a significant reduction in heterogeneity for both H3K27me3 fluorescence intensity and nuclear volume in the MPZ and PSM following TSA or Gar treatment (Fig. 6e-f). Notably, the MPZ showed a more pronounced decrease in nuclear volume variance uponTSA or Gar exposure compared to the PSM (Fig. 6f). Importantly, live imaging revealed that embryos treated with TSA or Gar experienced significant developmental delays, particularly in body axis elongation speed (Fig g-f), underscoring the critical role of chromatin remodeling-mediated collective cell movement in orchestrating zebrafish body axis elongation. These findings collectively establish a mechanistic link between cell fate decision and collective cell movement mediated by chromatin remodeling during zebrafish tailbud. Our results highlight the importance of epigenetic regulation in coordinating cell fate and tissue-scale morphogenetic processes during embryogenesis.

## DISCUSSION

The present findings illuminate how chromatin regulation and nuclear lamina organization link epigenetic states to tissue-level dynamics during zebrafish tailbud morphogenesis. Rather than focusing solely on nuclear volume, the data emphasize dynamic nuclear architecture—including chromatin condensation and lamina integrity—as a means for cells to coordinate the collective movements required for posterior body axis elongation. By integrating in vivo observations in the tailbud with in vitro epithelial monolayer experiments and epigenetic analyses, this study demonstrates that chromatin remodeling can fine-tune nuclear deformability, altering tissue fluidity and fate decisions among progenitor cells.

These results move beyond the view of isolated genetic or signaling pathways, shifting the focus to epigenetic modifications and nuclear framework that converge to guide tissue mechanics. Chromatin remodeling and nuclear lamina–mediated changes collectively drive transitions between fluid-like and jammed states, underlining how subcellular architecture can dynamically regulate cell motility. This new angle extends prior investigations, which often examined nuclear stiffness or volume in highly crowded tissues^15^, by revealing that chromatin- and lamina-driven remodeling exerts a more pervasive influence on morphogenesis than previously appreciated.

Experimental manipulations, such as suppressing chromatin modifications or altering Lamin A/C expression, highlight the importance of nuclear deformability in sustaining active cell rearrangements. Elevated Lamin A/C correlates with increased nuclear rigidity and diminished motility, paralleling features of a jammed state^34,35^. These observations fit into a broader framework wherein nuclear architecture—modulated by chromatin condensation and lamina composition—feeds back on cytoskeletal organization and cell–cell contacts, ultimately shaping tissue mechanics. Consequently, epigenetic changes can act on relatively short developmental timescales, continuously adjusting the nuclear structure to meet emerging morphogenetic demands.

By explicitly incorporating chromatin regulation and lamina integrity as nuclear drivers, this work refines existing cellular jamming models, such as vertex and Voronoi models, which have considered cell shape and density as drivers of jamming transitions^36–38^. The present framework suggests that nuclear “active fluctuation” levels—set by chromatin- and lamina-based remodeling—modulate cell shape changes and rearrangements. This perspective better accommodates developmental contexts in which nuclear plasticity under different genetic and epigenetic states is crucial for processes like tailbud elongation.

Although the data underscores how chromatin remodeling and lamina organization can influence tissue fluidity, several limitations remain. First, the current approach did not fully address the timeframe of epigenetic feedback on cytoskeletal restructuring or cell–cell adhesion. Time-resolved imaging of both chromatin states and mechanical properties would clarify how subcellular architecture evolves during critical developmental windows. Second, while the mechanism was validated in zebrafish tailbuds and cultured epithelia, it remains to be determined whether similar subcellular regulation governs morphogenesis in other vertebrates or in more complex three-dimensional organoid models.

Collectively, these findings emphasize that chromatin remodeling and the nuclear lamina converge to regulate nuclear deformability, linking epigenetic states and embryonic tissue mechanics. Demonstrating that chromatin remodeling-driven regulation of nuclear volume can orchestrate transitions between fluid-like and solid-like states redefines the conceptual framework of cellular jamming and underscores how epigenetic dynamics drive vertebrate morphogenesis as well as progenitor cell fate decisions. Future work integrating genetic, biophysical, and modeling approaches will be crucial for dissecting how epigenetic modifications operate in real time to guide subcellular structure and function. This line of inquiry promises deeper insight into how developing organisms coordinate their complex forms through the continuous interplay of chromatin, nuclear lamina, and cellular mechanics.

## MATERIALS AND METHODS

### Zebrafish Husbandry, Immunofluorescence Staining

Zebrafish (Danio rerio) were purchased from the National Zebrafish Resource Center in China and maintained under standard conditions, as previously described. All procedures were approved by the Institutional Animal Care and Use Committee (IACUC) and followed institutional guidelines. Wild-type embryos were collected from natural spawning. Embryos were raised in E3 medium at 28.5°C in a 14:10 hour light-dark cycle. Experiments were performed on 10-somite stage embryos. For imaging, embryos were manually dechorionated and mounted in 0.8% low-melting-point agarose (in E3 medium).

Dechorionated zebrafish embryos were fixed in 4% paraformaldehyde (PFA) at 4°C overnight. Fixed embryos were washed in phosphate-buffered saline (PBS) and permeabilized with 0.5% Triton X-100/PBS for 30 minutes at room temperature, followed by blocking in 5% bovine serum albumin (BSA) for 1 hour. Primary antibody incubation was performed overnight at 4°C in 1% BSA/PBS. After washing, the embryos were incubated with secondary antibodies for 2 hours at room temperature. Nuclei were stained with DAPI (1:500). Chromatin status was assessed using H3K27me3 (Cell Signaling Technology, 9733S; 1:400), and the secondary antibody used was Alexa Fluor 555- conjugated (Thermo Fisher, 1:500). Embryos were imaged using a Zeiss LSM 980 confocal microscope with a 40X water immersion objective, with a Z-step size of 0.4 μm.

### Cell Culture

MDCK (Madin-Darby Canine Kidney) cells were used as the model epithelial cell line for all experiments. Cells were cultured in DMEM (Dulbecco’s Modified Eagle Medium) supplemented with 10% fetal bovine serum (FBS) and 1% penicillin-streptomycin. Cultures were maintained at 37°C in a 5% CO₂ atmosphere.

### Inhibitor Treatments

To modulate chromatin condensation, two inhibitors were used: TSA (a histone deacetylase inhibitor, MCE, Cat# HY-15144, at a concentration of 20 nM) and Garcinol (a histone acetyltransferase inhibitor, MCE, Cat# HY-107569, at a concentration of 10 μM). TSA increases histone acetylation, typically leading to a more relaxed chromatin structure, while Garcinol reduces histone acetylation, promoting chromatin condensation. Both inhibitors were dissolved in DMSO and applied to MDCK cells at the specified concentrations for 24 hours. Control cells were treated with an equivalent volume of DMSO. After the treatment, cells were fixed in 4% paraformaldehyde and subsequently subjected to immunofluorescence staining to analyze chromatin state and nuclear volume fluctuations.

### Immunofluorescence and confocal microscopy

Cells were fixed in 4% paraformaldehyde (PFA) for 10 minutes, permeabilized with 0.2% Triton X- 100 in PBS for 10 minutes, and blocked in 5% bovine serum albumin (BSA) for 1 hour at room temperature. The samples were then incubated overnight at 4°C with primary antibodies in 1% BSA/0.2% Triton X-100/PBS, followed by washing in PBS and incubation with secondary antibodies in 1% BSA/0.2% Triton X-100/PBS for 2 hours at room temperature. The primary antibodies used were: HP1 (Abcam, ab109028; 1:400), H3K9me3 (Cell Signaling Technology, 13969S; 1:400), H3K27me3 (Cell Signaling Technology, 9733S; 1:400), Lamin A/C (Abcam, ab238303; 1:400). The secondary antibodies used were: Donkey anti-Rabbit IgG (H+L) Highly Cross-Adsorbed Secondary Antibody, Alexa Fluor™ 555 (Thermo Fisher, A-31572; 1:200), Goat anti-Mouse IgG (H+L) Cross-Adsorbed Secondary Antibody, Alexa Fluor™ 488 (Thermo Fisher, A-11001; 1:200), Donkey anti-Mouse IgG (H+L) Highly Cross-Adsorbed Secondary Antibody, Alexa Fluor™ 647 (Thermo Fisher, A-31571; 1:200). Nuclei were stained with DAPI (1:500). All fluorescence images were captured using a laser scanning confocal microscope (Zeiss LSM 980) with a ×40/1.1 C-Apochromat water-immersion objective (Zeiss), with a Z-step size of 0.4 μm. Images were acquired by sequential scanning at room temperature. The same imaging settings were applied for all samples in each experiment.

### Electron Microscopy Observation

To observe nuclear structure in detail, cells were imaged using transmission electron microscopy (TEM). Cells were fixed with 2.5% glutaraldehyde for 1 hour, followed by post-fixation with 1% osmium tetroxide. After dehydration through a graded ethanol series, samples were embedded in Epon 812 resin. Ultrathin sections (70 nm) were stained with uranyl acetate and lead citrate, and imaged using a JEOL JEM-1400 transmission electron microscope. TEM images were analyzed to assess nuclear morphology and chromatin condensation.

### Image processing

For MDCK brightfield images, cell area segmentation and calculations were performed using Cellpose 2.0 or Cellpose 3.0 in combination with ImageJ. For DAPI-stained fixed cells and zebrafish nuclei, as well as long-term fluorescently labeled nuclei in MDCK or zebrafish cells, Imaris 9.9 or arivis Pro 4.2.1 were used for 3D nuclear reconstruction, trajectory tracking, and data analysis. For long-term brightfield imaging of cell monolayers, PIVlab 3.0, a MATLAB-based software, was employed for velocity statistics. For live-cell brightfield imaging of zebrafish, ImageJ was utilized for length tracing and measurements.

### Live Imaging

Live-cell imaging was performed using an Andor spinning disk confocal microscope equipped with a 40X oil immersion objective to observe the nuclei of GFP-NLS-labeled MDCK cells. The experiment included pharmacological treatments with TSA and Garcinol (Gar). The drugs were applied to the cells three hours prior to imaging to allow for sufficient treatment effects. Following the treatment, live-cell imaging commenced, capturing the dynamics of nuclear volume fluctuations.

The total duration of the experiment was approximately 12 hours, with images captured every 10 minutes. The Z-axis step size was set to 0.35 μm to ensure a detailed 3D reconstruction of the cell nuclei.

### Drug Washout Experiment

For the drug washout experiment, a Leica microscope equipped with a 20X objective was used, with images captured every 5 minutes. Three hours after the initial imaging, drug treatment (TSA or Garcinol) was administered. The treatment lasted for over 18 hours, after which the drugs were washed out. Cells were then monitored for an additional over 24 hours to assess changes in cellular behavior.

### Plasmid Construction and Lentiviral Packaging

The LaminA/C plasmid was constructed using the PLVX-EGFP-IRES-PURO vector. The LaminA/C coding sequence was inserted into the multiple cloning site of the vector through restriction enzyme digestion and ligation, and the integrity of the construct was confirmed by sequencing. The plasmid was purified using a Qiagen plasmid midi kit. The GFP-NLS plasmid was constructed by cloning the nuclear localization signal (NLS) sequence into the pEGFP-C1 vector (Clontech). The NLS sequence was PCR amplified and inserted into the vector using restriction enzyme digestion and ligation. The successful construction of the plasmid was verified by Sanger sequencing and purified using a Qiagen plasmid midi kit.

The LaminA/C plasmid was transfected using lentiviral packaging. Lentiviral particles were produced by co-transfecting 293T cells with the PLVX-EGFP-IRES-PURO-LaminA transfer plasmid, the psPAX2 packaging plasmid, and the pMD2.G envelope plasmid (Addgene), using Lipofectamine 3000 (Invitrogen). Forty-eight hours post-transfection, the viral supernatant was collected, filtered, and used to infect MDCK cells in the presence of 8 μg/ml polybrene (Sigma). Infected cells were selected with 2 μg/ml puromycin (InvivoGen) for 5 days, and stable cell lines were confirmed by Western blot and fluorescence microscopy for LaminA expression.

For GFP-NLS transfection, Promega’s FuGENE 4K transfection reagent was used. One microgram of GFP-NLS plasmid DNA was transfected into MDCK cells at 60-70% confluency. The cells were incubated for 24-48 hours before further analysis or imaging to assess nuclear localization.

### Microinjection and Live-Cell Imaging

RNA for H2B-RFP can be made using in vitro transcription from the plasmids pCS-H2B- mRFP1( Plasmid #53745) purchased from addgene. These plasmids should be linearized with NotI, purified using a Qiagen nucleotide removal spin column, and used as template for in vitro transcription with the mMESSAGE mMACHINE™ SP6 Transcription Kit (Thermo, AM1340). In vitro transcribed RNA can be purified using a MEGAclear™ Transcription Clean-Up Kit (Thermo, AM1908) and quantitated on a NanoDrop spectrophotometer. RNA integrity can also be assessed using gel electrophoresis.

Zebrafish embryos at the 1-4 cell stage were injected with 3-4 nl of mRNA solution using a WPI PV830 microinjector (WPI, USA) under a stereomicroscope. Embryos were placed in 1% agarose-coated injection plates, and mRNA was injected either into the yolk or directly into the blastomere. Injections were completed within 20 minutes post-fertilization to ensure consistency across all embryos. After injection, embryos were cultured in E3 medium at 28.5°C until reaching the 8-somite stage. Successful injections were verified by observing RFP fluorescence in the nuclei under a fluorescence microscope (Olympus, Japan) 6 hours post-injection.

For live-cell imaging, embryos at the 8-somite stage were manually dechorionated using fine forceps and mounted in 0.8% low-melting-point agarose (in E3 medium) in a glass-bottom imaging dish. The dish was placed on the stage of a Zeiss LSM 980 confocal microscope equipped with a 40X/1.1 C-Apochromat water-immersion objective. Embryos were imaged in a temperature-controlled environment maintained at 28.5°C. Z-stacks were captured with a Z-step size of 0.4 μm, covering the entire nucleus. Time-lapse imaging was performed every 10-12 minutes for 3-4 hours to capture nuclear dynamics during development. Images were acquired using ZEN software (Zeiss).

### ScATAC-seq Data Sources and Integration

The zebrafish scATAC-seq dataset originates from the publicly available NCBI repository (GSE243256, https://www.ncbi.nlm.nih.gov/geo/query/acc.cgi?acc=GSE243256). The corresponding BED file containing all sample data, GSE243256_ZEPA.All.sample.bed.gz, was downloaded for analysis. Metadata associated with the scATAC-seq dataset was obtained from https://doi.org/10.6084/m9.figshare.25465477.v1. From the “Cell annotation in ZEPA.xlsx” file, we extracted cell barcode information corresponding to the 14hpf developmental stage. These 14hpf- specific cell barcodes were subsequently filtered from GSE243256_ZEPA.All.sample.bed.gz. Based on the experimental batch information, the zebrafish 14hpf scATAC-seq data were segregated and saved as ZEPABatch1.bed.gz, ZEPABatch3.bed.gz, and ZEPABatch5.bed.gz. The publicly available zebrafish 14hpf scRNA-seq dataset (seurat_object_hpf14.rds) was sourced from the http:// tome.gs.washington.edu/. Additionally, zebrafish motif data, comprising 551 motifs, was derived from the CIS-BP database (https://cisbp.ccbr.utoronto.ca/).

### Zebrafish scATAC-seq Analysis

The scATAC-seq analysis was conducted using the R package ArchR (v1.0.2). Initially, zebrafish annotation files were constructed via createGeneAnnotation and createGenomeAnnotation. The zebrafish 14hpf scATAC-seq dataset was then processed into Arrow files using createArrowFiles. Subsequently, an ArchR analysis object was established through ArchRProject to facilitate downstream analyses. To detect and eliminate doublets, addDoubletScores was employed. Cells were filtered based on stringent quality control criteria, retaining only those with TSSEnrichment ≥ 5 and nFrags ≥ 2000. Dimensional reduction was performed on the ArchRProject using addIterativeLSI with an iterative LSI approach (corCutOff = 0.75), followed by addHarmony to correct for batch effects across samples. A t-SNE embedding was generated within the ArchRProject via addTSNE (perplexity = 200, dimsToUse = 1:100, maxIterations = 1000, force = T). Clustering analysis was performed using addClusters, yielding 32 distinct clusters (C1–C32) at a resolution of 1. The addGeneScoreMatrix function computed gene scores for each gene across individual cells. To achieve cross-platform integration of scATAC-seq and scRNA-seq data (seurat_object_hpf14.rds), addGeneIntegrationMatrix was utilized. The scRNA-seq annotations were subsequently mapped onto the scATAC-seq clusters, with both clusters C4 and C23 identified as tailbud mesoderm.

We extracted tailbud mesoderm cells and constructed a new ArchRProject. Subsequently, we designated C4_Tailbud mesoderm as MPZ and C23_Tailbud mesoderm as PSM. To identify characteristic peaks for each specified cell group, getMarkerFeatures was employed with PSM as the reference background (useMatrix = “PeakMatrix”). The addMotifAnnotations function was then used to annotate the newly constructed ArchRProject with motif information present within the detected peaks. Enrichment analysis was conducted using peakAnnoEnrichment, which annotated the characteristic peaks with their corresponding motifs and identified significantly different motifs based on the criteria (cutOff = “FDR <= 0.05 & Log2FC >= 2”). A heatmap was generated to visualize motif scores for differentially enriched motifs between MPZ and PSM. To highlight key motifs, plotBrowserTrack was utilized to showcase the distribution of critical transcription factors, including tbx16, tbx6, tbxta, sox2, zeb1b, and snai1a. To further investigate lineage dynamics, addTrajectory was applied to analyze the developmental trajectories of MPZ and PSM. Finally, plotTrajectory was used to visualize the trajectory maps of MPZ and PSM while displaying the gene scores of key motifs along the trajectories.

Levene’s test was employed to assess the statistical significance of variance differences in peak scores and motif scores between MPZ and PSM. A p-value < 0.05 indicated a significant disparity in variance between the two groups. Similarly, the Wilcoxon test was utilized to evaluate the median differences in peak scores and motif scores between MPZ and PSM. A p-value < 0.05 signified a statistically significant difference in medians between the two groups.

### Data Analysis

Data processing was performed using MATLAB, and T-tests and visualizations were conducted with built-in R algorithms. Variance and kurtosis values were calculated using MATLAB’s built-in std and kurtosis functions. For calculating ΔV_nucleus, nuclear data for each trajectory were extracted in MATLAB, and theoretical nuclear volumes at corresponding time points were obtained using linear fitting with the polyfit function. The difference between the actual nuclear volume and the theoretical nuclear volume at each time point was defined as ΔV_nucleus for that trajectory.

The probability distribution function for 1/Area was generated using the cdfplot function. For ΔV_nucleus-velocity and 1/Area-shape index plots, the x-axis parameters were divided into equal intervals, and the mean of all corresponding y-values was calculated for each x-point. Outlier points with fewer than 15–20 data points were removed.

For heatmaps, a 25x25 grid was used, with the average velocity of all data points within each grid block mapped to the color bar. Heatmaps were generated using the heatmap function in MATLAB.

### Effective disk model with size fluctuations

In this model, nuclei are considered as 2D disks whose radii are fluctuating. The assembly of nuclei is then modeled by a semi-grand ensemble of polydisperse disks, as previously studied in [PHYS. REV. X 8, 031050 (2018), PRE 99, 012106 (2019)]. The interactions between nuclei are not explicitly known. Without loss of generality, they are assumed to experience a modified Lennard-Jones (LJ) potential. This specific choice of interaction potential is not essential to our main conclusion, i.e., without changing other parameters such as the volume fraction, increasing the radii fluctuations can fludize the system and cause a jamming-to-unjamming transition. In order to model the nuclei radii fluctuations, the semi-grand ensemble is introduced, which imposes a chemical potential 𝜇(λ_i_) to the *i*-th particle with the radius λ_i_. In such a setup, both particle positions and radii are treated as degrees of freedom, which evolve during the equilibrium procedure. In particular, the disks perform “breathing-like” motion, mimicking the nuclei volume fluctuations. The stable states are obtained by minimizing the total interaction and chemical potential energy, and then their diffusivity at is examined to determine if the configurations are jammed.

In detail, the total potential energy is:

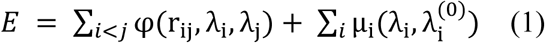

where φ(r_ij_, λ_i_, λ_j_) is the modified LJ potential expressed as:

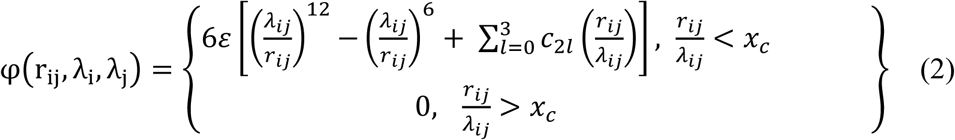

where 𝜀 is the energy parameter, λ_*i*_ is the *i*^𝑡ℎ^ particle’s radius or effective size, λ_*i*𝑗_ = λ_*i*_ + λ_𝑗_ and 𝑐_2𝑙_ is computed at conditions that the first three derivatives of φ with respect to 𝑟_*i*𝑗_ continuously vanish at the dimensionless cut-off 𝑥_𝑐_ = 2.0. The chemical potential μ_i_(λ_i_, λ_i_^0^) is given by:

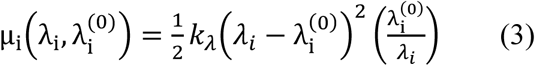

where 𝑘_λ_ is the stiffness parameter, and λ_i_^(0)^ = 0.5 is prefixed before the dynamical simulation. Importantly, it has been shown that the size polydispersity (the dimensionless variance of the radius distribution) is proportional to 1/ 𝑘_λ_ [PHYS. REV. X 8, 031050 (2018), PRE 99, 012106 (2019)], and thus we use the parameter 𝛿𝑅 = 1/𝑘_λ_ as a measure of the radius fluctuation.

The mechanical equilibrium states of the model are generated in the following way. We begin to equilibrate the system by initially setting the number density 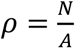 (A is the area) at 0.5 and the temperature at T = 1.0. The system is then minimized using FIRE algorithm [Physical review letters 97.17 (2006): 170201] subjected to the potential 𝐸 in Equation (1). This minimization is performed while fixing the pressure 𝑝 = 1.0 using Berendsen barostat with a time constant 𝜏_𝐵𝑒𝑟_ = 10.0. States are considered at mechanical equilibrium when the derivative of the potential over interparticle distance (i.e., the typical net force) is less than 10^−12^. Upon the convergence of minimization process, we perform an additional step to minimize the LJ potential part only in Equation (1), by freezing the effective radius λ_i_. We perform this final equilibration until the system reaches the mechanically stable packing configuration (total potential energy per particle < 10^−16^). This additional step guarantees the mechanical equilibrium of the final configuration.

Once mechanical equilibrium states are obtained, we perform overdamped dynamical simulations. For comparison, we compare two types of dynamics: with and without “breathing-like” motion described by the second term μ_i_ in Eq. (1). The disk motion follows Newtonian dynamics simulated by the standard numerical Verlet-integration method, with a timestep 𝛿𝑡 = 0.005. The effective mechanical force 𝐹_*i*_ is computed using the relation 𝐹_*i*_ = −∇_*i*_𝐸, with or without the second term μ_i_ in Eq. (1) included. The mass, length and time are expressed in term of the LJ-reduced units. The dynamical order parameter to distinguish jammed (𝐷_𝑒ff_ > 10^−3^) and unjammed states is the self-diffusivity 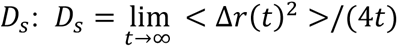, where <Δ𝑟(𝑡)^2^> is the mean-squared displacement.

#### Self-propelled Voronoi model with shape fluctuations

The model describes each cell as a Voronoi tessellation domain whose vertices are treated as the degrees of freedoms. The cell center and the Voronoi vertices are related via the Delaunay-Voronoi duality. The intercellular interaction among N cells is controlled by their perimeter and area. The total mechanical energy of the system is expressed as:

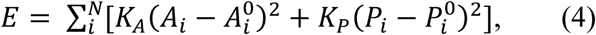

where 𝐴_*i*_ and 𝑃_*i*_ are the area and perimeter of the *i*-th cell, 𝐴_*i*_^0^ and 𝑃_*i*_^0^ are pre-fixed reference parameters, 𝐾_𝐴_ and 𝐾_𝑃_ are the area and perimeter modulus characterizing how hard it is to change the cell’s area and perimeter. The total energy is a result of a competition between targeted perimeter (𝑃*_i_*^0^) and area (𝐴_*i*_^0^), giving rise to the so-called effective shape index: 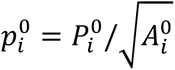. This quantity controls the preferred shape cells, and the occurrence of the jamming-unjamming transition. In previous studies, 𝑝_*i*_^0^ is generally a constant. In contrast, in the present work, 𝑝_*i*_^0^ of each cell is drawn randomly from a Gaussian distribution characterized by pre-selected mean 𝑝_0_ and standard deviation 𝛿𝑝_0_.

The equation of motion is given by overdamped dynamics with the self-propelled active force:

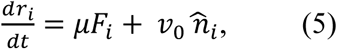

where 𝜇 is the inverse of the frictional drag, 𝐹_*i*_ is the effective mechanical force 𝐹_*i*_ = −∇_*i*_𝐸 with E given by Eq. (4), 𝑣_0_ = 0.1 is the activity. The unit 𝑛^_*i*_ follows random rotational diffusion,

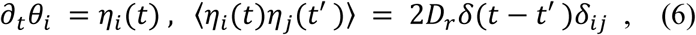

where 𝐷_𝑟_ = 1 determines the memory of the stochastic noise.

The dynamical order parameter to distinguish jammed and unjammed states is similarly the self-diffusivity 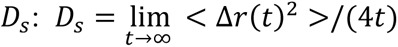. In practice, the scaled diffusivity 𝐷_𝑒ff_ = 𝐷_𝑠_/𝐷_0_, where 𝐷_0_ = 𝑣_0_^2^/(2𝐷_𝑟_), serves as a parameter to differentiate unjammed (𝐷_𝑒ff_ > 10^−3^) and jammed (𝐷_𝑒ff_ < 10^−3^) states.

## ACKNOWLEDGMENTS

This study was supported by the National Natural Science Foundation of China (NSFC) (12222201, 82273500, 82372750, 12332019, U20A20390), the National Key R&D Program of China (2023YFC2507000), Fundamental Research Funds for the Central Universities (ZG140S1971, KG16228501) and the CAMS Innovation Fund for Medical Sciences (2023-I2M-3-003 and 2022-I2M- 1-008).

