## Supplementary figures and images for "Chromatin remodeling integrates vertebrate body axis elongation and cell fate determination"

### Supplemental Figure

Supplement Fig 1

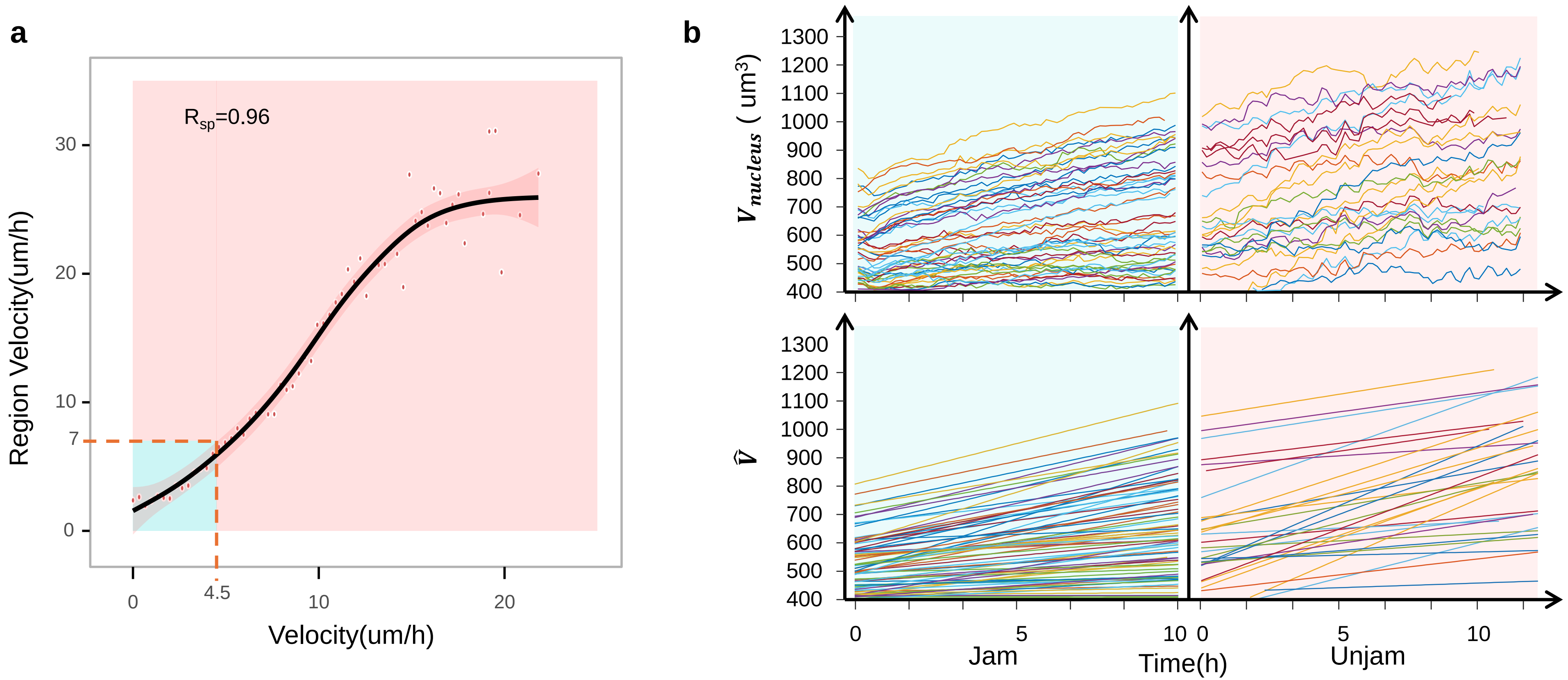

Supplement Fig 2

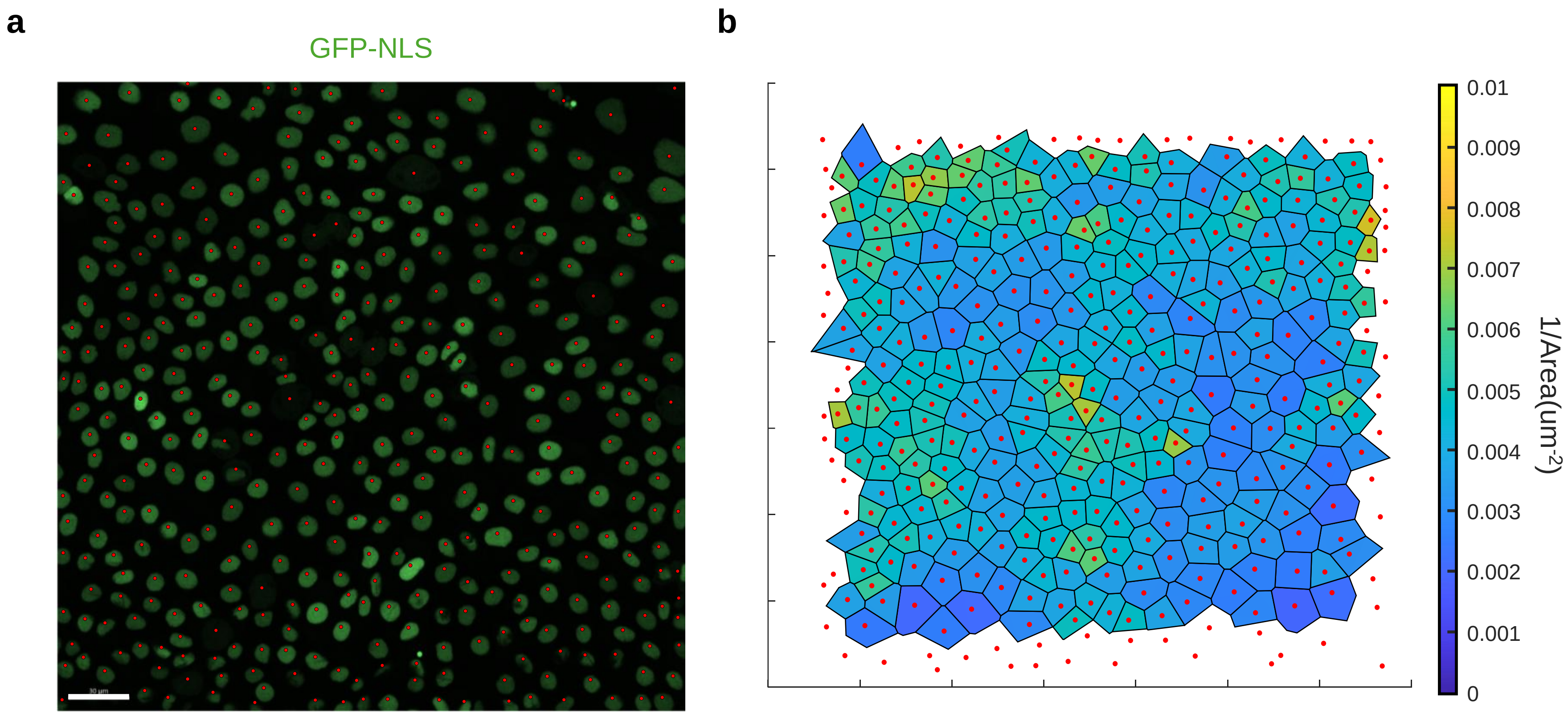

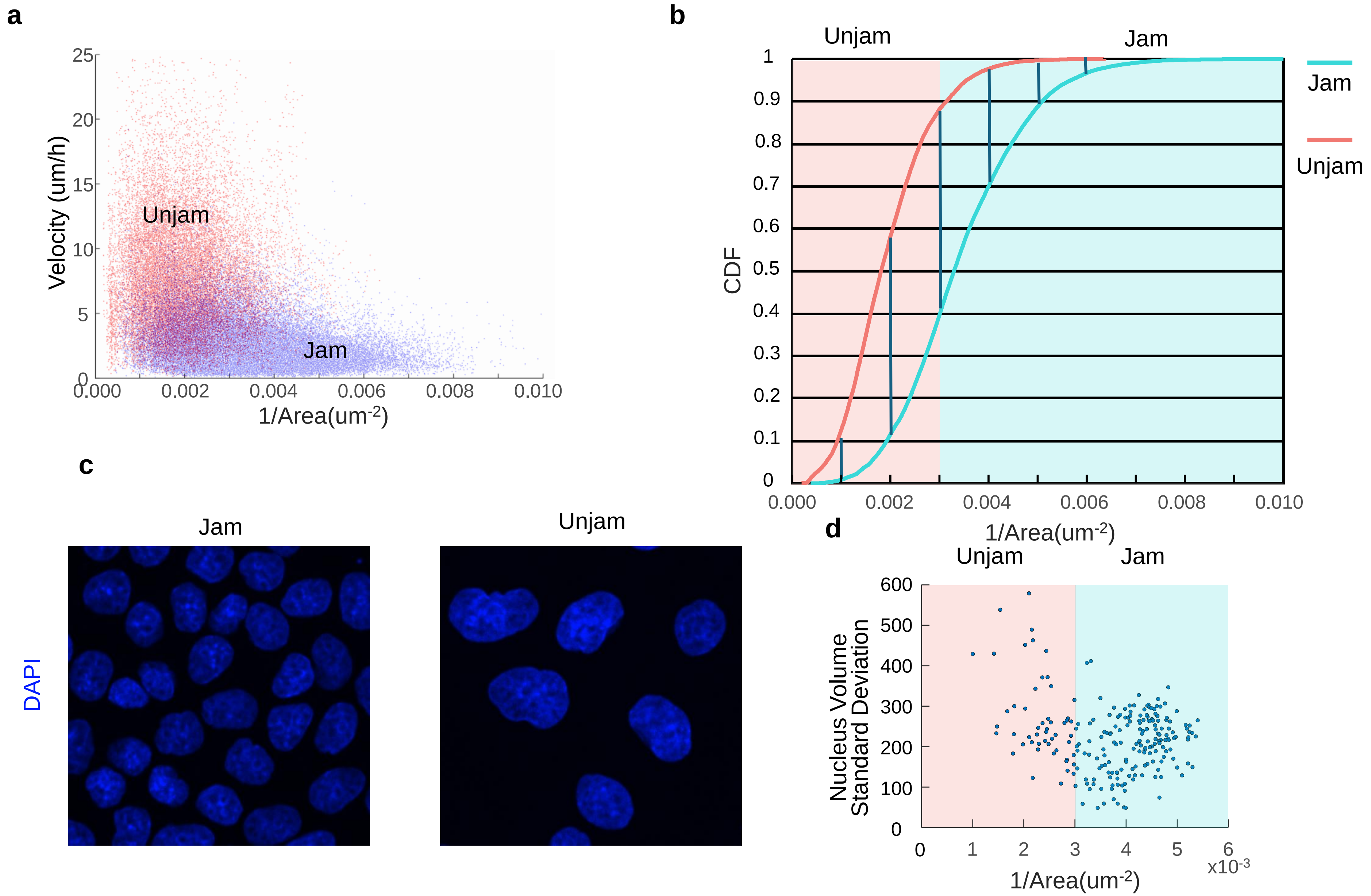

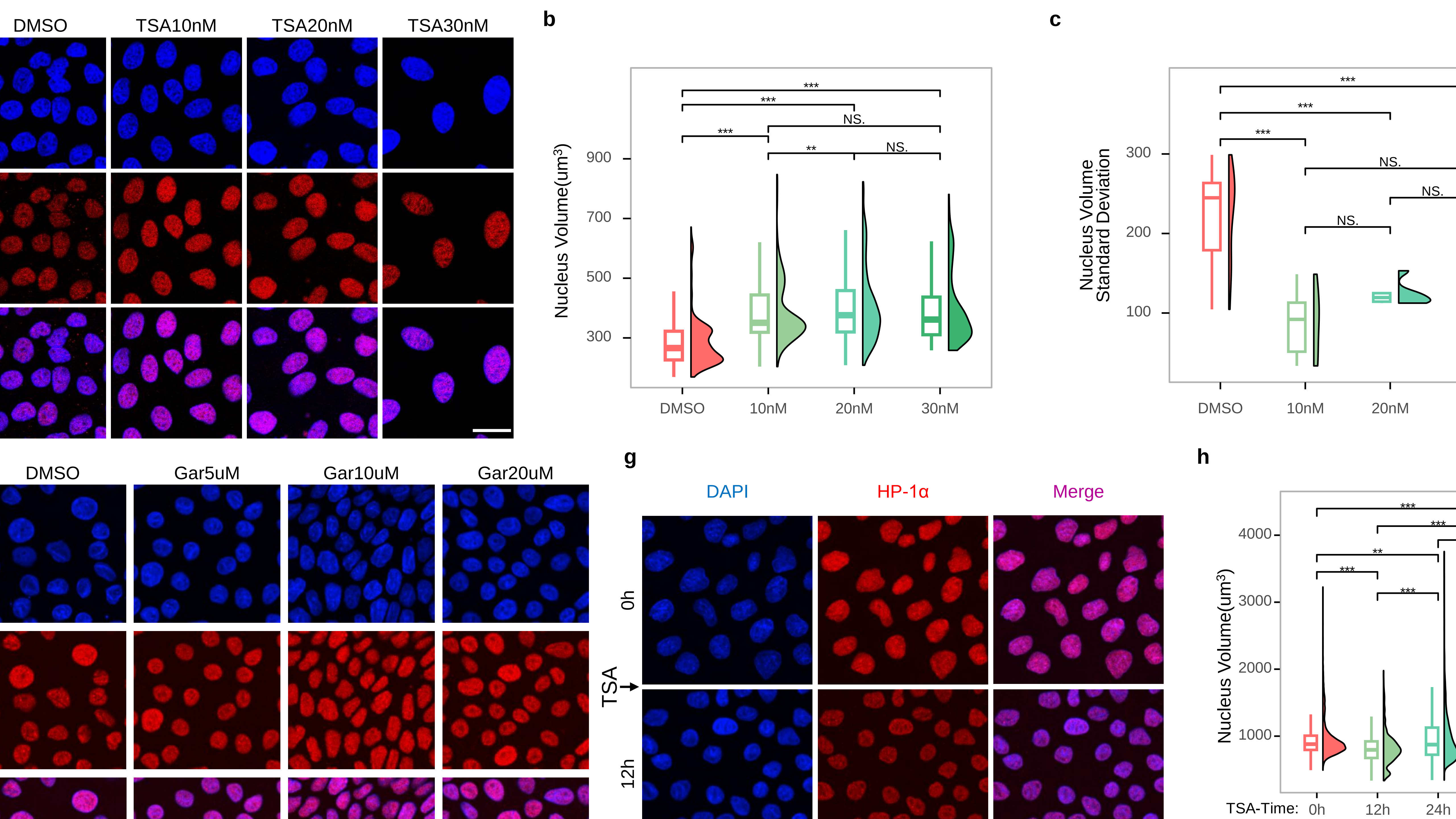

Supplement Fig 5

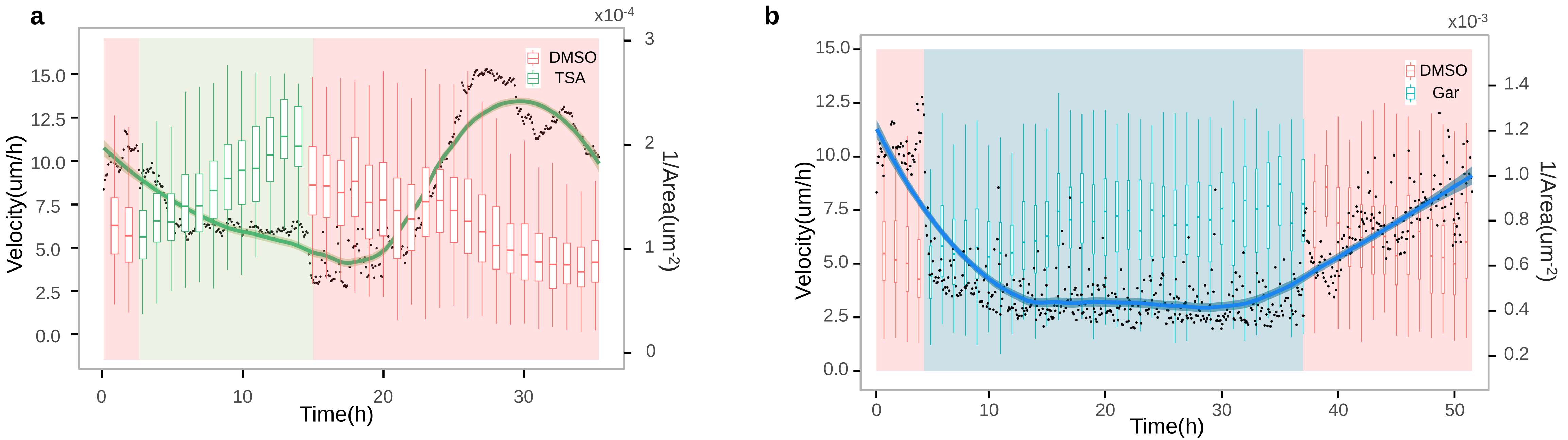
